# Evolution of generalists by phenotypic plasticity

**DOI:** 10.1101/761288

**Authors:** David T. Fraebel, Karna Gowda, Madhav Mani, Seppe Kuehn

**Affiliations:** Department of Physics. University of Illinois at Urbana-Champaign. Urbana, IL 61801, USA; Center for the Physics of Living Cells. University of Illinois at Urbana-Champaign. Urbana, IL 61801, USA; Carl R. Woese Institute for Genomic Biology. University of Illinois at Urbana-Champaign. Urbana, IL 61801, USA; Department of Molecular Biosciences, Northwestern University, Evanston, IL 60208, USA; NSF-Simons Center for Quantitative Biology, Northwestern University, Evanston, IL 60208, USA; Department of Engineering Sciences and Applied Mathematics, Northwestern University, Evanston, IL 60208, USA

## Abstract

Adapting organisms face a tension between specializing their phenotypes for certain ecological tasks or developing generalist strategies which permit persistence in multiple environmental conditions. Understanding when and how generalists or specialists evolve is therefore an important question in evolutionary dynamics. Here we study the evolution of bacterial range expansions by selecting *Escherichia coli* for faster migration through porous media containing one of four different sugars supporting growth and chemotaxis. We find that selection in any one sugar drives the evolution of faster migration in all sugars relative to the ancestral strain. Measurements of growth and motility of all evolved lineages in all nutrient conditions reveal that the ubiquitous evolution of fast migration arises via phenotypic plasticity. Phenotypic plasticity permits evolved strains to exploit distinct phenotypic strategies to achieve fast migration in each environment irrespective of the environment in which they were evolved. Our work suggests that plasticity plays an important role in the evolution of generalist phenotypes.

## Introduction

Organisms in nature often encounter varied environments throughout their lifetime, each with its own demands on the phenotype. Therefore, the ability of a single genotype to thrive under different environmental conditions can be essential for a lineage’s chance of long-term persistence. Organisms that can thrive in varied environments are called generalists. It is thought that generalists evolve in fluctuating environments, while fixed environments select for specialists [1]. In its native environment the specialist is expected to have enhanced fitness relative to the generalist, but exhibit reduced fitness in other environments [2].

However, the results of experimental evolution studies show remarkably diverse outcomes. While trade-offs are sometimes observed [3], they are far from universal: studies with a constant selection environment (which are expected to produce specialists) often produce a mixture of specialists and generalists across replicate lineages [4]. Additionally, trade-offs observed in experimental evolution are sometimes asymmetric: studies employing multiple selection conditions may produce specialists in one condition and generalists in another [5, 6]. Other studies find that a majority of evolved lines are generalists, even with selection performed in a constant environment [7, 8].

A key question which emerges from these studies is understanding the limits and mechanisms of the evolution of phenotypic generality. Experiments suggest that generalists will only show enhanced fitness in environments that are sufficiently similar to their selection condition [9, 10]. While this notion is intuitive, there are distinct avenues of phenotypic adaptation that could give rise to generalist phenotypes. An experimentally supported interrogation of the conditions under which generalists evolve, the environmental limits of their generality, as well as its underlying genotypic and phenotypic mechanisms would shed light on how populations deploy phenotypic variation during evolution to adapt [11].

To address this, we selected *Escherichia coli* for faster migration through porous media with one of four different sugars as a carbon source and chemoattractant. We observed the evolution of generalists: selection for fast migration in any one sugar resulted in fast migration in all sugars. Because migration depends on both growth and motility, we measured these phenotypes for strains from each evolutionary history in all four nutrient conditions to investigate the evolution of migration rate generality. We found that the environment determined the phenotype more than the evolutionary history: strains from all selection conditions exhibited a distinct growth and motility phenotype for each assay condition irrespective of their evolutionary history. That is, all strains observed in a particular environment showed similar adaptation of growth and motility regardless of their selection condition. Because the evolved strains in this study exhibit different phenotypes depending on their environment, and because those phenotypes are associated with faster migration, we conclude that the nutrient generality of migration rate evolution was achieved through phenotypic plasticity [12].

## Results

### Repeated selection enhances migration of bacterial populations through soft agar

*E. coli* inoculated into low viscosity agar deplete nutrients locally as cells swim and divide in the porous, three-dimensional environment. This depletion establishes a nutrient gradient which drives chemotaxis outwards and subsequent growth of the population [13, 14]. The result is a macroscopic colony that expands radially from the site of inoculation at a speed determined by growth, motility and chemotaxis of its constituent cells [15]. We performed time-lapse imaging on these colonies and observed an initial growth phase followed by radial expansion at a constant rate. See supplemental information (SI) for images of expanded colonies in each condition during selection.

We performed experimental evolution by selecting *E. coli* for faster migration through soft agar. After allowing a population to expand for 24 hours, we selected a small population of cells from its outermost edge and used them to inoculate a new low viscosity agar plate (Figure 1a). This process of expansion and selection was repeated for ten rounds. From a single, ancestral strain (MG1655-motile) we performed selection for faster migration in M63 minimal medium with 0.2 % w/v agar and one of four different sugars as carbon sources at 1 mM concentration: mannose, melibiose, N-acetylglucosamine and galactose. In each condition, the sugar served as the sole energy source and chemoattractant for the expanding population. We chose these four sugars because we believed that the diverse set of genetic architectures involved in their chemotaxis, import and metabolism could lead to diverse opportunities for genomic evolution and resultant phenotypic adaptation in each condition. All four sugars traverse the outer membrane through *OmpF*, but mannose and N-acetylglucosamine use phosphotransferase systems to cross the inner membrane while melibiose and galactose rely on cation symporters *MelB* and *GalP* [10, 16, 17]. Once inside the cell, catabolism of N-acetylglucosamine and galactose is regulated by repression from *NagC* and *GalR/GalS* [18, 19]. On the other hand, melibiose catabolism is regulated by activation from *MelR* [20] and mannose catabolism does not have a specific inducer, requiring only the global regulator cyclic AMP receptor protein (CRP) to signal glucose starvation [21].

**Figure 1:**
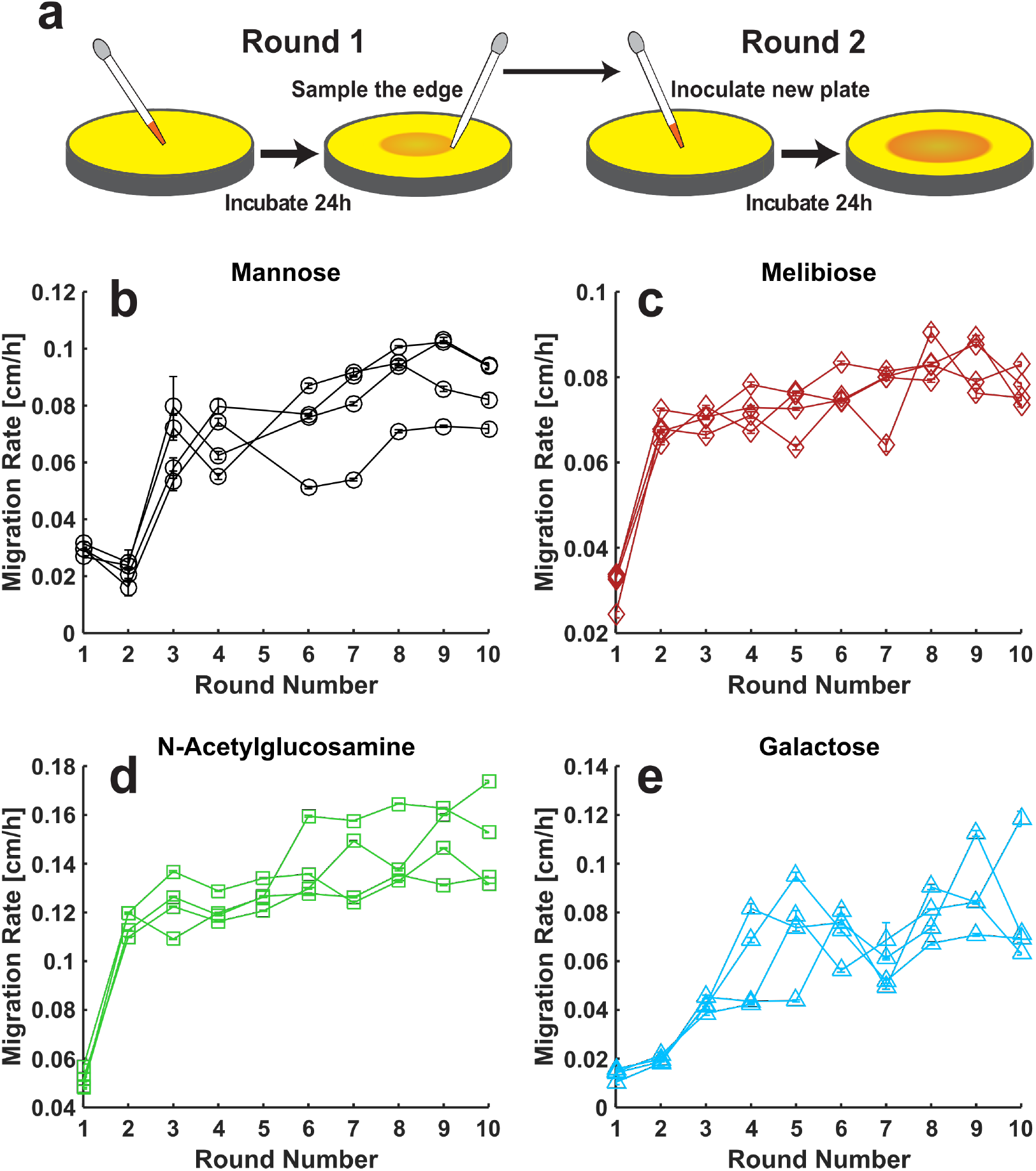
Repeated selection enhances *E. coli* migration through soft agar in four nutrient conditions. **(a)** Schematic of migration selection procedure. Motile *E. coli* are inoculated into the bulk of a soft agar plate containing growth medium. Nutrient consumption and chemotaxis drive the growing population to expand radially across the plate at a constant migration rate. After a fixed interval, cells are sampled from the edge of the expanding colony and used to inoculate a new plate. **(b-e)** Migration rates as a function of round of selection for experiments conducted in 0.2 % w/v agar plates containing M63 minimal medium with one of four different carbon sources at 1 mM concentration: mannose, melibiose, N-acetylglucosamine and galactose. Four replicate experiments were carried out to 10 rounds in each condition. Migration rates are measured by time-lapse imaging of expanding colonies followed by linear fit of colony radius versus time; error bars are 95 % confidence intervals on fitted rates. No rates are reported for round 5 in mannose due to failure of the imaging device. See methods for details of selection procedure and migration rate measurement.

Across all four conditions, we observed a dramatic enhancement of migration rates due to selection. Migration rates of all lineages in melibiose and N-acetylglucosamine more than doubled after just one round of selection (Figure 1c,d), while similar improvement in mannose and galactose was achieved after two rounds of selection (Figure 1b,e). In all four conditions, migration rates continue to improve in subsequent rounds, albeit modestly compared to the initial increase. In round 10, migration rates had increased almost threefold in mannose and N-acetylglucosamine, 2.5-fold in melibiose, and nearly 6-fold in galactose.

### Selection for fast migration in one sugar results in fast migration in all sugars

We wondered whether migration rate evolution was specific to each selection condition or whether strains evolved in one sugar would migrate fast relative to the ancestor in other sugars. Therefore, we measured the migration rates of evolved strains from each evolutionary history in all nutrient conditions. Specifically, four independent replicate lineages from each selection condition (a total of 16 strains) had their migration rates measured in four different assay conditions: mannose, melibiose, N-acetylglucosamine and galactose. Surprisingly, nearly all strains exhibited enhanced migration across all conditions relative to the founding strain in that condition (Figure 2). For example, a strain evolved for fast migration in N-acetylglucosamine (squares, Figure 2) exhibited faster migration than the ancestral strain in all three of the other sugars. Migration rates of non-native strains (that is, populations being assayed in a condition different from their selection condition) were nearly always well in excess of the ancestor’s rate, often comparable to or even exceeding the natively-evolved populations’ migration rates. The only exceptions to this statement are two melibiose evolved strains migrating in galactose which were on average only 1.58 ± 0.09 times faster than the founder, compared to all other stains being 4.49 ± 1.33 times faster than the ancestor in this condition, mean ± standard deviation (Figure 2d). The non-native populations migrate quickly despite having no evolutionary history in that nutrient condition. The objective of this study is to understand how selection in one nutrient environment generically gives rise to fast migration in all nutrient environments.

**Figure 2:**
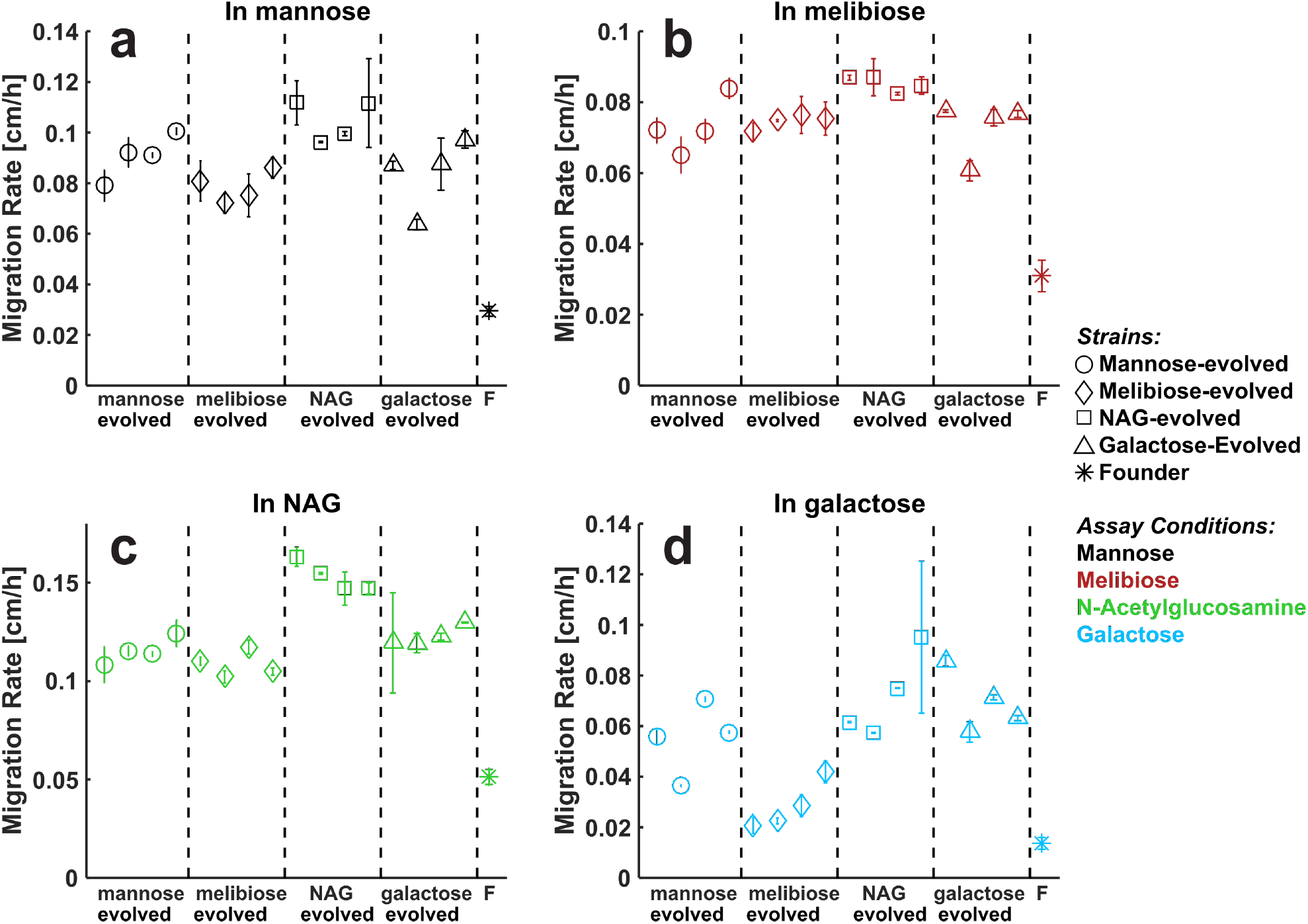
Nutrient generality of migration rate evolution. The 16 strains isolated after 10 rounds of selection (four from each nutrient condition, Figure 1b-e) were assayed for enhanced migration rate in each of the four nutrient conditions used for the selection experiments. Migration rates of evolved strains are presented as mean ± standard deviation of two replicate plates for each strain in each condition. Marker shapes denote selection condition, marker colors denote assay condition (legend). Due to failure of the imaging device, migration rates of the rightmost two galactose-evolved strains in galactose **(d)** are reproduced from round 10 of the selection experiments (Figure 1e). Migration rates of the founding strain (F) in each condition are presented as mean ± standard deviation of the four migration rates measured in each condition during round 1 of the selection experiments (Figure 1b-e).

To establish the limits of this generality, we first measured the migration rates of the ancestor as well as two evolved strains from each selection condition in seven additional nutrient environments beyond the initial four: arabinose, dextrose, fructose, lactose, maltose, rhamnose and sorbitol. Again, we found that nearly all the strains showed enhanced migration rates relative to the ancestral strain across all these conditions (SI). The evolved strains typically migrated 1.5 to 3 fold faster than the ancestor. We concluded that the nutrient generality of migration rate selection extends to many different carbon sources within the regime of M63 with 0.2 % w/v agar and 1 mM sugar. In contrast, migration rates of all 16 evolved strains presented in Figure 2 did not exhibit fast migration in lysogeny broth rich medium (LB) and instead exhibited a drop in migration rate relative to the ancestor. For comparison, we also measured three independently-evolved strains isolated after 10 rounds of selection in LB (SI) and found that they migrate around 50 % faster than the ancestor. So while repeated selection still enhances migration in LB, the generality of the strains evolved in minimal medium does not extend to this rich medium where chemotaxis and growth are driven by amino acids [13].

### What makes a generalist?

We next set out to understand how phenotypes evolved under selection for fast migration in one sugar give rise to fast migration in other sugars. Population-level migration through soft agar depends on both growth and motility of individual cells. Therefore, selection for faster migration can be driven by enhancements to growth rate, chemotactic response and undirected motility [15, 22]. For example, increases in running speed or tumble frequency are known to drive faster migration through soft agar [23]. What phenotypic changes permit a lineage selected on one sugar to migrate more rapidly than the ancestral population in another sugar (Figure 2)?

To understand why strains from distinct evolutionary histories (selection conditions) show enhanced migration across different environments (assay conditions), we must consider the growth and motility of each evolved strain in each assay condition. For the present discussion we consider growth and motility as abstract traits but our hypotheses do not depend number or identity of the traits considered. In this framework, we propose that there are three main possibilities for how the generality of fast migration could evolve (Figure 3).

**Figure 3:**
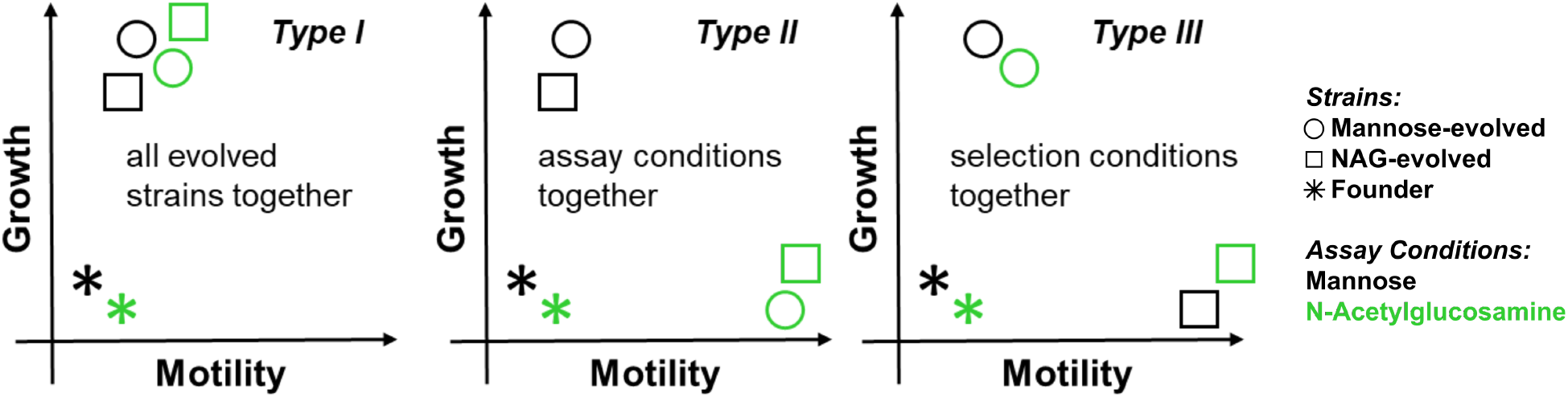
Scheme to characterize the phenotypic basis of migration rate generality. We present three distinct possibilities for the underlying phenotypic basis of the nutrient generality of migration rate adaptation presented in Figure 2. *Type I: Universality* All evolved strains show the same adaptation from founder across all assay conditions. There is no separation of evolved phenotypes in the two dimensional phenotypic space of motility and growth. *Type II: Plasticity* Evolved strains exhibit a flexible adapted state. Evolved strains show a characteristic adaptation for each assay condition, regardless of their evolutionary history. Phenotypes separate by assay condition (marker color). *Type III: Degeneracy* Evolved strains display a characteristic phenotype for each evolutionary history, regardless of assay condition. Phenotypes separate by selection condition (marker shape).

#### Type I: Universal adaptation

In this scenario neither the selection condition nor the assay condition has an impact on the evolved phenotype. Instead, there is a single phenotype conferring fast migration through 0.2 % w/v agar and 1 mM sugar, irrespective of any differences in import, metabolism and chemotactic affinity between different sugars. Evolved strains could achieve this phenotype across all selection conditions. Generality would then be achieved as long as evolved strains exhibited the adapted phenotype across different assay conditions. For example, the evolved strains could have achieved a growth rate adaptation that does not depend on the specific sugar, allowing them to grow fast (and thereby migrate fast) in all 1 mM sugar assay conditions. Or, they could exhibit a change in run-tumble statistics which confer a migration rate advantage in soft agar. However, *Type I* generality need not be this simple. The ideal phenotype could be a particular combination of growth and motility enhancements, corresponding to a distinct direction away from founder in the two dimensional space of phenotypes. The key feature of *Type I* generality is that there is no separation of evolved phenotypes by either selection condition or assay condition. Instead there is a single, universal, phenotype which is achieved by all evolved strains in all conditions (Figure 3).

#### Type II: Phenotypic plasticity

In each sugar, there exists a distinct growth/motility phenotype conferring fast migration. The adaptive value of growth or motility can easily depend on the assay condition. For example, in a condition that supports slow growth of the ancestral strain, increases in growth may confer a greater advantage compared to motility. These differences could also arise at the molecular level due to differences in import, metabolism and chemotactic affinity for each sugar. In this scenario, generality would emerge if evolved strains exhibit the different adapted phenotypes for each assay condition, irrespective of their selection condition. This would mean that selection acts to put evolved populations in an adapted, plastic state. Once in this state, cells could adapt their phenotype to the particular balance of growth and motility needed to enhance migration in different environments. We call this mechanism for evolving generalists plasticity because it requires that the same genotype (evolved strain) exhibit distinct phenotypes (growth/motility) in different nutrient conditions. In this situation, a strain’s phenotype is determined more by its assay condition than selection condition. Graphically, this would mean that the evolved phenotypes separate by assay condition (Figure 3). For example, if evolved strains across different evolutionary histories showed enhanced growth in mannose but enhanced motility in N-acetylglucosamine, we would conclude that the nutrient generality of migration rate evolution was achieved through phenotypic plasticity.

#### Type III: Degeneracy

In this scenario there is a degenerate set of distinct phenotypes which all confer faster migration on the evolved strains in all assay conditions. Suppose that there is a distinct method of adaptation associated with each selection condition and evolved populations exhibit these different adaptations in all assay conditions. Generality would be possible in this case as long as different growth and motility phenotypes could produce similar migration rates in a particular environment. In this sense the evolved phenotypes would be degenerate at the level of migration rate. Graphically, this would correspond to the case where evolved phenotypes separate by selection condition (Figure 3). For example, if mannose-evolved strains enhance growth in all environments, while strains evolved in N-acetylglucosamine enhance motility across all environments, we would classify the observed migration rate generality as *Type III*.

### Measured phenotypes suggest generality evolved by plasticity

Within this framework we sought to establish which type of generality was responsible for the generalist adaptation we observed in selection for faster migration. To accomplish this goal we set out to quantify the phenotypes relevant for bacterial migration for each evolved strain in all four environments. Three processes define the population-level migration rate: bacterial diffusion (with cellular diffusion constant *D*_*b*_), chemotactic response (with strength *χ*) and growth (with growth rate *k*_*g*_) [24]. In the simplest case, where no chemotaxis is present, the classic result from Fisher shows that a diffusing and growing population will form a traveling wave with speed 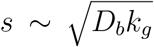. Allowing chemotaxis to bias the motion of individual cells in response to local nutrient gradients enhances the migration rate. In the situation where chemotaxis and growth are driven by a single nutrient, the migration rate scales as 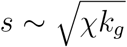, where *χ* itself depends on the bacterial diffusion constant and the strength of the cell’s response to a gradient [15],[Cremer *et al.* In revision, 2019 (personal communication)]. Quantifying bacterial diffusion constant and growth in high-throughput is feasible, but methods for reliably quantifying *χ* are low typically low throughput [25] or prone to large errors [26].

Therefore, to gain insight into the mechanism responsible for the migration rate generality we observed, we measured the growth and diffusion of the ancestor as well as the 16 evolved strains presented in Figure 2 in each of the four environments. These experiments were conducted in liquid minimal medium identical to the plates used for migration assays, but without agar. Motility was measured by performing high-throughput single-cell tracking on populations of cells with phase contrast microscopy [27] and diffusion constants (*D*_*b*_) were inferred from the slope of the mean squared displacement across individual trajectories (Figure 4a). We note that measured diffusion constants in liquid closely match diffusion the 0.2 % w/v agar condition where our selection experiment took place [15]. Growth was measured by monitoring the optical density of well-mixed liquid cultures and maximum growth rates (*k*_*g*_) were fitted during the exponential phase of the growth curve (Figure 4b).

**Figure 4:**
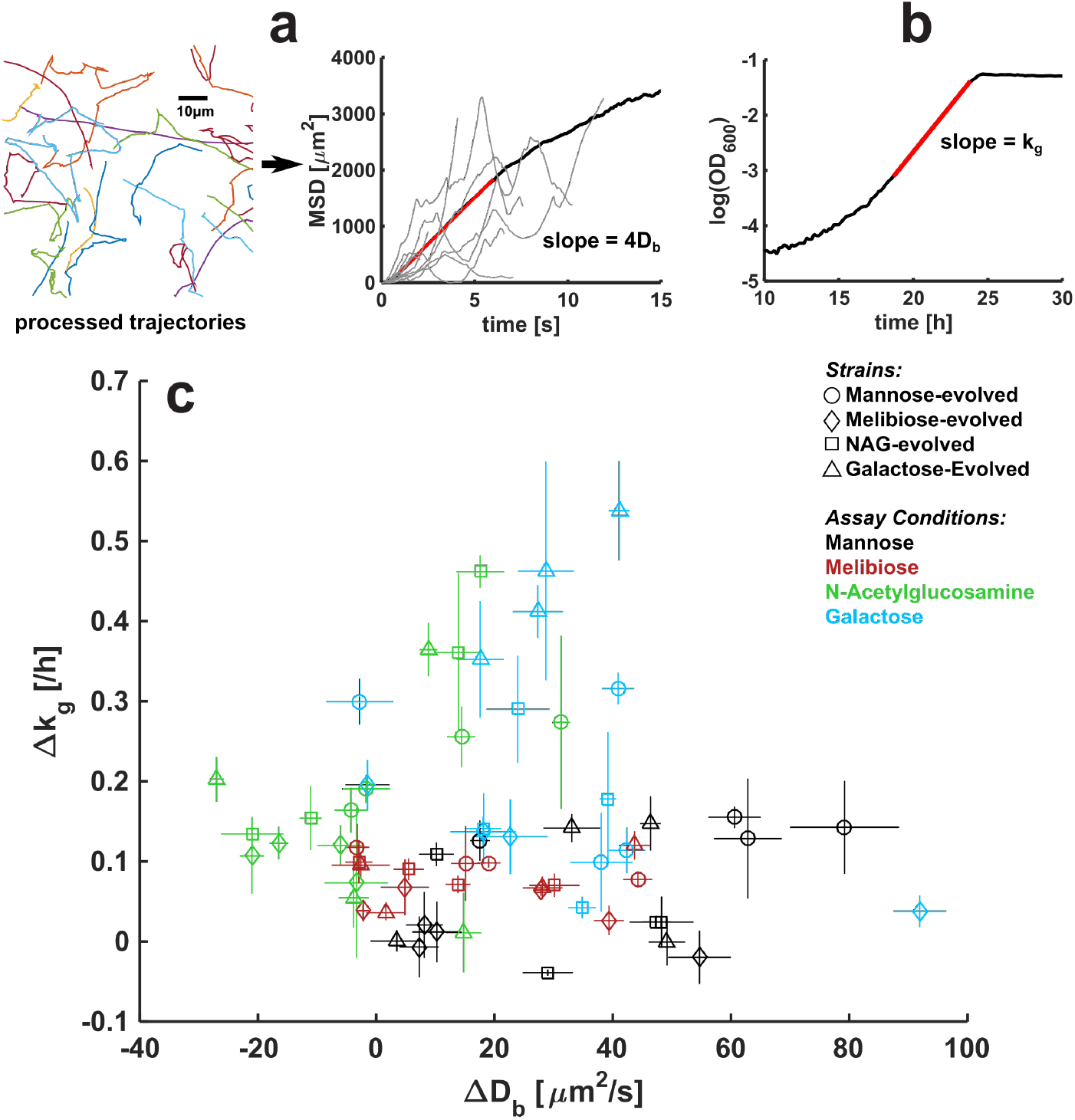
Phenotypic adaptation suggests plasticity. **(a)** Scheme for measuring bacterial diffusion constants. For each strain in each condition, thousands of individuals were recorded swimming on glass slides (7856 ± 5623, mean ± standard deviation trajectories per experiment). Videos were automatically processed into trajectories and squared displacement was calculated for each cell at each frame. Examples of single-cell squared displacement traces given in light gray. Population-level mean squared displacement (MSD, black line) was calculated by averaging over single-cell traces at each frame. *D*_*b*_ was inferred from a linear fit to the MSD vs time (red line). **(b)** Maximum growth rates were measured by continuously measuring the optical density of well-mixed liquid cultures. *k*_*g*_ was fitted from the slope of *log*(*OD*_600_) versus time and averaged over replicate wells. See methods for details of both experiments. **(c)** Motility and growth adaptation in liquid media of all 16 evolved strains presented in Figure 2 in four different assay conditions. The founder has a diffusion constant of 54 ± 19, 69 ± 13, 61 ± 10 and 56 ± 0.6 μm^2^/s in mannose, melibiose, NAG and galactose respectively, mean ± standard deviation of two replicate experiments. The founder has a maximum growth rate of 0.18 ± 0.05, 0.29 ± 0.01, 0.30 ± 0.05 and 0.16 ± 0.08 h^−1^ in mannose, melibiose, NAG and galactose respectively, mean ± standard deviation of ten replicate wells spread over two independent plates. The evolved phenotypes have been subtracted by these values to present adaptation of diffusion constant (Δ*D*_*b*_) and adaptation of maximum growth rate (Δ*k*_*g*_). Error bars on evolved diffusion constants are 95 % confidence intervals on fitted slopes from mean squared displacement versus time. Error bars on evolved growth rates are standard deviations across five replicate wells.

Since we are interested in how the evolved strains differ from the ancestor in each condition, we chose to subtract the evolved phenotypes by the ancestor’s *k*_*g*_ or *D*_*b*_ in the same condition. Thus, we will investigate the distribution of evolved phenotypes in the two dimensional space of Δ*k*_*g*_ and Δ*D*_*b*_ to distinguish between the three types of generality presented in Figure 3. The results of these measurements are shown in Figure 4c.

A quick inspection of the evolved phenotypes hints at phenotypic plasticity. The data seems to roughly separate by assay condition (color, Figure 4c). For example, strains measured in mannose typically have a larger enhancement to diffusion but a more modest enhancement to growth when compared to strains measured in galactose. We next sought a statistical measurement of this effect to evaluate any significant separation of evolved phenotypes by selection condition and/or assay condition. To achieve this, we performed analysis of variance (ANOVA) on the following linear mixed effects model.

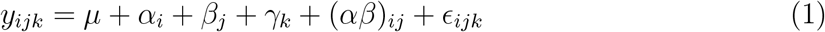

In this model, we want to understand a response variable *y*_*ijk*_ (Δ*k*_*g*_ or Δ*D*_*b*_) in terms of deviations from the global mean *μ* attributable to different groupings of the data. *α*_*i*_ is a fixed effect due to selection condition. *β*_*j*_ is a fixed effect due to assay condition. *γ*_*k*_ is a random effect for each of the 16 evolved strains. (*αβ*)_*ij*_ is an interaction term included to allow for unique effects due to particular combinations of assay and selection conditions. For example, if some strains grow particularly well in the condition they were selected in compared to the other strains, this would manifest itself as a significant interaction term. Lastly, *ϵ*_*ijk*_ is a noise term in the form of normally distributed random disturbances.

The ANOVA allows us to determine which groupings, if any, have significant differences between their levels. Performing the ANOVA on Δ*D*_*b*_, we found that assay condition is the only grouping with a significant (*p* < 0.05) F-statistic (Table 1). That is, we cannot reject the null hypothesis that all the *β*_*j*_ are zero. We conclude that there is a significant difference between the assay conditions in the adaptation of motility. To investigate which assay conditions show significant departures from the global mean, and in which direction, we did *post-hoc* testing via a bootstrapping approach. This revealed that strains measured in mannose and galactose were significantly above the global mean in Δ*D*_*b*_, while strains measured in N-acetylglucosamine were significantly below the mean and strains in melibiose showed no significant change in *D*_*b*_ relative to the global mean (SI).

**Table 1:** Assay condition drives differences in diffusion constant adaptation. Summary statistics for ANOVA on the model presented in Equation 1 using Δ*D*_*b*_ as the response variable. The F-statistic describes the ratio of between-group variability to within-group variability. Here, we only found a significant (*p* < 0.05) F-statistic for assay condition.

| Source of Variance | F | p-value |
| --- | --- | --- |
| Selection Condition ($\alpha_i$) | 0.84 | 0.50 |
| Assay Condition ($\beta_j$) | 11.95 | $1.41 * 10^{-5}$ |
| Strain ($\gamma_k$) | 1.81 | 0.08 |
| Selection Condition x Assay Condition ($(\alpha\beta)_{ij}$) | 0.64 | 0.76 |

Bacterial diffusion constants are reasonably approximated by the run speed and duration: 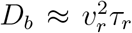. Therefore, changes in diffusion constants can be achieved through changes in run duration and/or speed. We wondered whether Δ*D*_*b*_ was achieved with a consistent microscopic strategy within each assay condition. Therefore, we investigated correlations between changes in run statistics and changes in diffusion constant across all evolved strains within each assay condition (SI). We found that Δ*D*_*b*_ was significantly correlated with a particular microscopic strategy for strains measured in each assay condition, regardless of their evolutionary history. For example, in mannose changes in *D*_*b*_ were correlated with changes in *τ_r_* but not *v_r_*. Conversely, in N-acetylglucosamine changes in diffusion constant are correlated with changes in both *v_r_* and *τ_r_*.

Similarly, ANOVA with Δ*k*_*g*_ as the response variable shows that by far the most significant effect is due to assay condition (largest F-statistic, Table 2). *Post-hoc* testing revealed significant departures from the mean for all four assay conditions: strains measured in N-acetylglucosamine and galactose have above-average growth rate adaptation, while strains measured in mannose and melibiose were below average (SI). We also find smaller but still significant (*p* < 0.05) effects due to selection condition and the interaction term. *Post-hoc* testing on the *α*_*i*_ revealed only one significant coefficient: melibiose-evolved strains exhibit a below average growth rate enhancement. For the interaction term, *post-hoc* showed only a few significant terms, most notably an increased growth rate adaptation for strains evolved in N-acetylglucosamine and galactose and assayed in their own selection conditions.

**Table 2:**
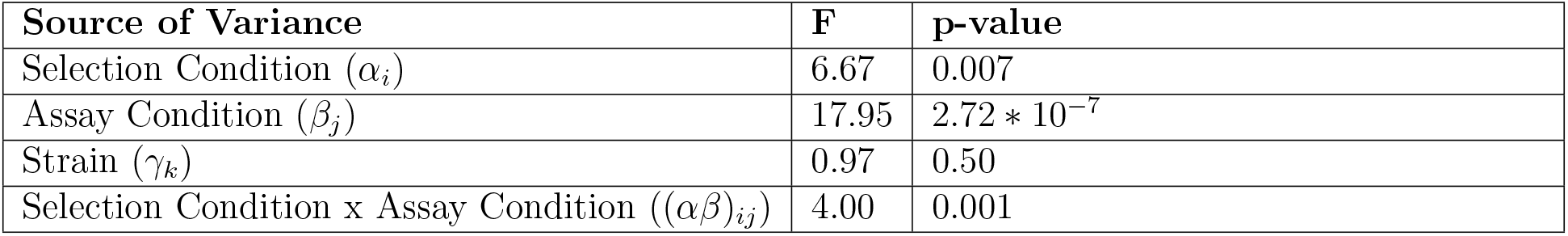
Assay condition is main driver of differences in growth adaptation. Summary statistics for ANOVA on the model presented in Equation 1 using Δ*k*_*g*_ as the response variable. The F-statistic describes the ratio of between-group variability to within-group variability. Here, we found significant F-statistics for selection condition, assay condition, and the interaction term between them. The largest F-statistic was associated with assay condition, indicating this predictor is the dominant source of variability in Δ*k*_*g*_.

Our statistical approach provides clear evidence that the evolved phenotypes presented in Figure 4c separate by assay condition. We conclude that within each of the four environments, evolved strains exhibit similar adaptation in growth and motility regardless of their evolutionary history. The result is consistent with *Type II* generality. Evolved strains across all selection conditions appear to have evolved phenotypic plasticity which allows them to enhance population-level migration by adapting cellular phenotypes to meet the unique demands of each nutrient condition.

### Mutations present in evolved strains cannot predict phenotypes in any condition

We next asked whether specific mutations present in strains evolved for faster migration could explain the migration, growth and motility phenotypes we observed. We performed whole genome sequencing on all 16 evolved strains and identified *de novo* mutations relative to the common ancestor present at a frequency of at least 20 %[28]. We identified 33 unique mutations occurring in diverse genes and pathways including biosynthesis of essential molecules, stress response and metabolite import (SI). We observed no single mutation common to all evolved strains nor any mutations common to all 4 evolved strains from any particular selection condition. However, 24 of the 33 mutations were present in more than one strain. By grouping together mutations occurring in the same gene as well as those in adjacent intergenic regions, we identified 21 unique mutational targets. 12 of those targets were affected in multiple strains, and of those, 8 were affected across strains with different selection conditions. Unlike previous studies [22], we did not observe mutations with an obvious interpretation in terms of their impact on evolved phenotypes. Therefore, we took a statistical approach to interpreting our sequencing data.

Having grouped mutations by their target, we created a mutation candidacy matrix (*X*) which describes the presence (1) or absence (0) of each observed mutation in each strain. We then asked whether the presence or absence of these mutations could predict the migration rate of each strain in each condition. To do this we performed linear regression using mutation candidacy as the predictor variables and the adaptation in either migration rate, growth rate, or diffusion constant as the dependent variable, e.g., 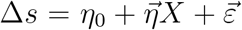, where *η*_0_ is an intercept, 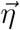 is a vector of regression coefficients for each mutation and 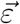 is a noise term with zero mean and variance *σ*^2^. We performed *L*_1_-regularized regression (LASSO) to avoid over fitting [29, 30] using leave-one-out cross validation to determine the best value of the regularization hyperparameter. For several assay conditions the LASSO procedure selected a model with only an intercept (e.g., giving 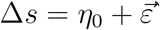). In most cases where a model with non-zero 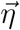 was selected, the improvement in the mean-squared error (estimated using cross validation) relative to a model with only an intercept was small.

These observations led us to undertake a numerical investigation of the performance of LASSO for our regressions (SI). Briefly, we constructed surrogate data sets which retained the statistical structure of the candidacy matrix *X*, but specified 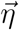 and the magnitude of *σ*^2^. We specified the number of nonzero entries in 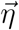, drawing these entries from a standard normal distribution. We constructed surrogate data sets using these synthetic 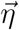, performed regressions and constructed a statistic which estimates the improvement in model fits (mean-squared error estimated using cross-validation) for the model selected by LASSO relative to a model with only an intercept (*η*_0_). We measured the quality of each model fit by making out-of-sample predictions on additional surrogate data. We found that in a high noise regime (*σ* > 1), the out of sample predictive power of the models tended to be poor even if the number of nonzero entries in 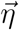 was small (i.e., if the number of mutations impacting the phenotype is small). These numerical experiments showed that our test statistic correlated positively with the out-of-sample predictive power of a model inferred by LASSO. We then computed our test statistic for regressions which predicted adaptation in migration rate, growth rate and diffusion constants from the mutation candidacy matrix and concluded that, for our data, mutations are highly unlikely to predict the measured phenotypic adaptation (out-of-sample *R*^2^ ≈ 0). The sole exception to this conclusion was the regression predicting migration rate adaptation in mannose.

## Discussion

The central finding of this study is the emergence of *Type II* generality when bacterial populations are selected for faster migration through a porous minimal medium environment. We found that distinct phenotypic strategies gave rise to fast migration and, remarkably, these strategies were determined more by the nutrient condition of the measurement than the evolutionary history of the strains being measured. We concluded that repeated selection in any condition drives fast migration in all conditions via distinct, plastic phenotypic responses to each nutrient condition.

The molecular mechanisms which give rise to strains evolved in different environments exhibiting similar phenotypes when assayed in the same environment are not yet clear. Our statistical analysis of the genetic variation observed in evolved strains shows that there is no simple genetic basis for this plastic adaptive response. Given what is known about sugar uptake and metabolism in *E. coli*, some of our observed mutations could be targeting the uptake or metabolism of multiple sugars. For example, *EnvZ* regulates expression of *ompF*, the outer membrane responsible for import of all four sugars [31, 10]. *NagA* is essential for metabolism of N-acetylglucosamine, but also has a role in regulating expression of the *nag* regulon, including *nagC* [18]. There is evidence that *NagC* is capable of repressing the mannose PTS system [18] and the galactose transporter [32]. Therefore, the mutations we observed in *nagA* for two galactose-evolved strains could have an impact in mannose, N-acetylglucosamine and galactose environments. Our observation of similar phenotypic outcomes arising in strains with distinct genetic variation is consistent with the idea that adaptation can be facilitated by mutations modifying weak regulatory interactions in the cell [33].

The mechanistic basis of how metabolism is coupled to motility remains unclear and elucidating this fully will be required to understand why evolved strains in some conditions exhibit large changes in both motility and growth (e.g. galactose). The fact that the evolved strains do not exhibit fast migration in rich media where amino acids are responsible for growth and chemotaxis is consistent with the idea that the plastic generalist adaptation is specific to sugars. Therefore the limits on generalist evolution due to plasticity in this system would appear to be defined by the chemical identity of the nutrient/chemoattractant. Given that we observe mutations in regulatory elements (e.g. *rssB, wzzE, envZ*) it is possible that the plasticity we observe in the evolved strains could be best understood at the regulatory level. In this case, the shared molecular-level features of the evolved strains which are responsible for the phenotypic outcomes in different nutrient conditions may be explained by gene expression measurements [34].

Since the seminal work of Baldwin and later Waddington [35, 36], a main focus of work on phenotypic plasticity has been to understand the relationship between plastic responses to environmental change and subsequent genetic adaptation [37, 38, 39]. Our results suggest another possible role of phenotypic plasticity in evolution, namely, that selection in one environment can potentiate new physiological responses to other environments resulting in the evolution of generalists even in homogeneous environmental conditions. Future work should focus on understanding why selection in one environment can elicit new phenotypic responses to other environments and the scope of phenotypic generality that can evolve by this mechanism.

## Methods

### Migration rate assay and selection experiment

Plates were prepared by autoclaving agar into laboratory-grade water, cooling to 55 °C, then adding sterile stock solutions of M63 media components and the desired carbon source on a heated stir plate. The media had a final agar concentration of 0.2 % w/v and a final sugar concentration of 1 mM. 22 mL of media was added to a 10 cm petri dish and allowed to gel before being wrapped with parafilm and stored at 4 °C until use. Plates were thermalized for 24 hours before use in the 30 °C environmental chamber where all migration experiments were conducted (Darwin Chambers).

All migration assays were initiated by growing 5 mL cultures of *E. coli* (strain MG1655-motile, Coli Genetic Stock Center (CGSC) #8237) overnight to saturation from frozen stock in 5 mL of liquid M63 with 30 mM of sugar matching the plate to be used. 10 μL of saturated culture was injected into the center of a soft agar plate. Time-lapse imaging was performed for 24 hours at 5 minute intervals on the expanding colonies using webcams (Logitech HD Pro Webcam C920) in a dark box with pulsed illumination provided by warm white LED strips (LEDMO SMD2835) around the periphery of each plate. Automated image analysis was used to extract migration rates as described below.

For selection experiments, eight 20 μL samples were removed from the outermost edge of the expanding colony after imaging. The sample was briefly vortexed and 10 μL was immediately injected into a fresh plate from the same batch to initiate the next round of selection. The remainder of the sample was preserved at −80 °C on 25 % glycerol. This process of imaging, sampling and inoculation was repeated until ten rounds of selection had been completed, at which point a final sample was taken and preserved.

### Image analysis

Webcam-acquired images were processed by custom written Matlab code. First, images were background subtracted by an image created from six early time points before growth had occurred. The colony’s center was determined by applying Canny edge detection and a circular Hough transform to an image near the end of the experiment. Next, radial intensity profiles were constructed for each image along a line outwards from the colony’s center. Local cell density is monotonic with pixel intensity. Location of the colony’s edge was determined by applying a threshold to the intensity profiles, and migration rate was determined by linear regression of the edge’s position versus time during the final 5 hours of expansion. Calibration was performed by imaging a test target to determine the number of pixels per centimeter.

### Growth rate measurement

Seed cultures were grown from frozen stocks in 5 mL of M63 with 1 mM sugar for 36 hours to saturation. Cultures were then diluted 1 : 1000 into a 48-well plate containing 1 mL of fresh media in each well. Optical density was measured in a plate reader (Tecan Infinite 200 or BMG Labtech CLARIOstar) by 600 nm absorbance every 10 minutes with 200RPM shaking between measurements. Maximum growth rates (*k*_*g*_) were acquired by linear fit of the linear portion *log*(*OD*) versus time (exponential growth phase) just before the roll-off to stationary phase. Fitting intervals were determined manually and were typically two to five hours in duration.

### Single-cell tracking

Glass slides and cover slips were cleaned by sonication in acetone followed by 1 M KOH, passivated with 2 mg mL^−1^ bovine serum albumin and rinsed with laboratory-grade water before use. Cultures were grown from frozen stocks in 5 mL of M63 with 1 mM sugar to early-mid exponential phase, *OD*600 ≈ 0.14. 5 μL of culture was added to the passivated region of a slide, covered with the passivated side of a cover slip, and the chamber was sealed with Devcon 5 Minute Epoxy. Videos of swimming cells were acquired for 30 or 60 seconds at 12 frames per second with a Point Grey model FL3-U3-32S2M-CS camera and a phase contrast microscope (Omano OM900-T inverted) at 10x magnification. Illumination was provided by a high-brightness white LED (LED Supply 07040-PW740-L) to avoid the 60 Hz flickering that observed with the stock halogen lamp. Experiments were performed in a 30 °C environmental chamber (Darwin Chambers). For each evolved strain in each condition, 5 videos were acquired from different sites on the glass slide. For the founder, this process was repeated for two independent replicate slides in each condition.

Videos were processed into trajectories and run-tumble classified using PyTaxis (https://github.com/tatyana-perlova/pytaxis) [27]. Briefly, this software segments videos to obtain cellular coordinates, links coordinates from each frame into trajectories and filters out trajectories of cells stuck to the glass slide as well as trajectories fewer than 20 frames. Subsequently, filtered trajectories were analyzed using custom Matlab code. First, we calculated each cell’s squared displacement at each frame. Next, we averaged over cells to obtain a mean squared displacement versus time trace for each strain in each condition. A five-second interval near the beginning of the trace was observed to be linear across datasets, indicating diffusive motion of the cells. The slope of this region was used to compute the diffusion constant (*D*_*b*_) after dividing a prefactor of 4, since swimming cells were confined to two dimensions in this experiment. After this interval, the traces become sub-linear due to the presence of cells with lower diffusivity making up these very long trajectories (Figure 4a). Computing the distribution of trajectory lengths confirmed that such cells constitute a minority of the total trajectories (SI).

### Analysis of variance and post-hoc testing

We performed 3-way ANOVA on the model presented in Equation 1 using the Matlab function anovan. We used either Δ*k*_*g*_ or Δ*D*_*b*_ as the response variable, and selection condition, assay condition and strain as predictor variables. Selection condition, assay condition and the interaction term between those two predictors were all treated as fixed effects. Strain was treated as a random effect nested into selection condition. This serves as a strain-specific noise term which accounts for the lineage to lineage stochastic variation in phenotypes within one selection condition.

ANOVA results were used to determine which predictors have significant differences in the response variable between their levels. For predictors that showed a significant F-statistic, we further investigated which coefficients showed significant departures from the global mean using a non-parametric bootstrapping approach. To do this, we first randomly permuted the response variable to create a randomly-labeled dataset. We then performed the ANOVA on the randomly-labeled data and computed the coefficients for the predictor being studied. We repeated the process of random permutation and ANOVA 10 000 times to construct a distribution of ANOVA coefficients from randomly-labeled datasets. Finally, we computed a p-value by considering the fraction of coefficients from shuffled data larger/smaller than the coefficients from ANOVA on the properly-labeled data. The intent of this approach is to highlight the probability that the particular arrangement of selection conditions, assay conditions and strains we observed in the Δ*D*_*b*_, Δ*k*_*g*_ phenotypic space (Figure 4c) could arise due to random chance. If an ANOVA coefficient from our data is unlikely given a distribution of coefficients from ANOVAs on randomly-labeled data, we conclude that the associated group has a meaningful departure from the global mean in whichever response variable is being investigated.

### Whole genome sequencing and analysis

Cultures of each evolved strain were grown from frozen stocks in 5 mL LB overnight to saturation. Genomic DNA was purified using the Qiagen DNeasy UltraClean Microbial Kit. Input DNA was quantified by Qubit and sequencing libraries were prepared using the NexteraXT kit from Illumina. Purified, amplified libraries were quantified by Qubit and Bionanalyzer and normalized with the bead-based method of the NexteraXT kit. Sequencing was performed on an Illumina MiSeq system following pooling, dilution and denaturation of the bead-normalized libraries according to MiSeq system specifications. Reads were demultiplexed and adapter trimmed with the onboard Illumina software. Sequence data for the ancestral strain was used from our previous work, where we sequenced the founder with an average depth of 553*x* by aggregating reads from four separate sequencing reactions [22].

Analysis was performed using the *breseq* computational pipeline (http://barricklab.org/twiki/bin/view/Lab/ToolsBacterialGenomeResequencing) in polymorphism mode. Breseq uses an empirical error model and a Bayesian variant caller to predict mutations. The software uses a threshold on the empirical error estimate (E-value) to call variants [40]. The value for this threshold used here was 0.01, and we report all mutations present in evolved populations at a frequency of 0.2 or above [40]. All other parameters were set to their default values. Reads were aligned to the MG1655 genome (INSDC U00096.3). We note that breseq is not well suited to predicting large structural variation. We excluded any mutations present in the founder from further analysis in order to consider novel genetic variation present in the evolved strains.

## Acknowledgements

D.F. and S.K. acknowledge support from the National Science Foundation Physics Frontiers Center Program (PHY 0822613 and PHY 1430124). KG acknowledges support from the James S. McDonnell Foundation Postdoctoral Fellowship Award (#220020499). M.M. acknowledges the Simons Foundation (597491, M.M.). M.M. is a Simons Foundation Investigator. We would like to thank Jun Song for advice in selecting the form of the ANOVA model, Tatyana Perlova for providing technical support and software for the single-cell tracking assays and Elizabeth Ujhelyi for assistance with the sequencing.

