## Supplemental Information for "Evolution of generalists by phenotypic plasticity"

### Numerical investigation of LASSO regression

We investigated the ability of $L_1$-regularized regression (LASSO) to predict the values of phenotypic parameters from mutational target candidacy using a numerical analysis of surrogate data. Our objective was to assess when the LASSO procedure could reliably detect true nonzero regression coefficients.

We generated surrogate data from a model of the form $\vec{Y} = \eta_0 + \vec{\eta}X + \vec{\varepsilon}$. We varied two control parameters to generate these data, $P$ and $\sigma$: $P$ is the number of randomly-selected elements in the regression coefficient vector $\vec{\eta}$ that are drawn from a standard normal distribution, with all other elements set to zero; $\sigma$ controls the magnitude of the noise term $\vec{\varepsilon}$, whose elements are drawn from a normal distribution with mean zero and variance $\sigma^2$. In order to simulate the structure of the true mutation candidacy matrix, a predictor matrix $X$ is generated by randomly shuffling along columns of the candidacy matrix, which preserves the number of observations ($N = 16$), the number of mutational targets (21), and the frequency of each mutation. Without loss of generality, we set the intercept $\eta_0 = 0$. From $\vec{\eta}$, $X$, and $\vec{\varepsilon}$, we obtain the surrogate response vector $\vec{Y}$. To generate additional data to be reserved for out-of-sample testing, we repeat the shuffling procedure to generate a new predictor matrix $X^{OOS}$ with $N = 16$ observations, and sampled a new noise vector $\vec{\varepsilon}^{OOS}$ to obtain response values $\vec{Y}^{OOS} = \vec{\eta}X^{OOS} + \vec{\varepsilon}^{OOS}$.

For each surrogate data set, we used LASSO regression (MATLAB R2017b) to fit a linear model of the form $\vec{Y} = \eta_0 + \vec{\eta}X + \vec{\varepsilon}$. The LASSO procedure generated a set of models over a range of values for the regularization hyperparameter $\lambda$. Leave-one-out cross-validation was used to estimate the model mean squared error (MSE) as a function of $\lambda$. Model selection was performed by choosing the value of $\lambda = \hat{\lambda}$ that minimized the cross-validated MSE. At $\lambda = \hat{\lambda}$, LASSO regression produces estimates for the model parameters: $\hat{\eta}_0$, $\hat{\vec{\eta}}$. Additionally, we inferred a model with only an intercept (i.e., $\vec{Y} = \eta_0$, which is the model resulting from the limit $\lambda \to \infty$). We then constructed a statistic, $M$ which measures the improvement of the model MSE for $\hat{\lambda}$ relative to the $\lambda \to \infty$ limit:

$$M = \frac{(MSE_{\lambda \to \infty} - MSE_{\hat{\lambda}})}{SE_{\hat{\lambda}}},$$

where $MSE_{\lambda \to \infty}$ is the MSE value of the model with only an intercept, $MSE_{\hat{\lambda}}$ is the MSE of the model inferred at $\hat{\lambda}$, and $SE_{\hat{\lambda}}$ is the estimated standard error of $MSE_{\hat{\lambda}}$ determined by cross-validation (Figure S1a). Model evaluation was performed by computing the coefficient of determination $R^2$ of the selected model applied to the out-of-sample predictor $X^{OOS}$ and response $\vec{Y}^{OOS}$:

$$R^2 = 1 - \frac{\sum_i (y_i^{OOS} - \hat{y}_i^{OOS})^2}{\sum_i (y_i^{OOS} - \bar{Y}^{OOS})^2},$$

where $i$ is an index over the 16 data points in the out-of-sample data set ($\vec{Y}^{OOS}$), $\hat{y}_i{}^{OOS} = \hat{\eta}_0 + \vec{\hat{\eta}} x_i^{OOS}$ and $\bar{Y}^{OOS} = \frac{1}{N}\sum_i y_i^{OOS}$ and $N = 16$ is the number of data points.

We performed the steps above for a grid of $P$ and $\sigma$ values ($P \in \{1, 2, 4, 8, 16\}$ and $\sigma \in \{0.1, 0.2, 0.4, 0.8, 1.6, 3.2\}$), generating $10^4$ instances of surrogate data for each $(P, \sigma)$ combination. We chose these values of $\sigma$ to sample both high and low noise regimes for the surrogate data. The average magnitude of the $P$ non-zero entries in $\vec{\eta}$ is unity, so $\sigma$ values range from much smaller (0.1) to much larger (3.2) than the regression coefficients. These two limits on $\sigma$ characterize low and high noise regimes respectively.

The results of these simulations are shown in Figure S1b-c. For each $(P, \sigma)$ combination the heat map shows median values of $M$ across the $10^4$ instances of surrogate data. The values of $M$ are highest in a low noise regime, corresponding to $P > 1$ and $\sigma$ sufficiently small. Median out-of-sample $R^2$ values are also highest ($> 0.5$) in this low noise regime, suggesting that high values of $M$ indicate situations where the LASSO procedure yields good inferences of the regression coefficients. To make the relationship between $M$ and out-of-sample $R^2$ explicit, we combined the values of $M$ and $R^2$ obtained in all surrogate data sets from all models ($n = 3 \times 10^5$ in total) and binned values by $M$ in intervals of 0.5. Within bins, we computed the quartiles (25th, 50th, 75th percentiles) of $R^2$, which we plot in Figure S1d-f. We observed that the relationship between $M$ and out-of-sample $R^2$ is indeed monotonically increasing, with an interquartile range that decreases as $M$ increases. Thus, as $M$ increases, it is increasingly likely that LASSO has correctly inferred the linear model.

We next performed the LASSO procedure on the experimental data with the true mutational target candidacy matrix $X$ for three different response variables: adaptation in migration rate ($\Delta s$), growth rate ($\Delta k_g$) and diffusion constant ($\Delta D_b$). Separate regressions were performed for all evolved strains in all four nutrient conditions. Using these regressions performed on the true data we then computed our statistic $M$, and these values are shown as colored vertical lines in Figure S1d-f. As in the surrogate data simulations, for each real data regression there are 16 observations, 21 mutational targets, and leave-one-out cross validation is used for determining $\hat{\lambda}$. In nearly all cases, the values of $M$ obtained in these regressions are near zero, and the numerical simulations indicate that an out-of-sample $R^2 \ll 0.5$ is likely. One exception is the regression on migration rate adaptation assayed in mannose, which yields $M = 6.4$. For this regression, the numerical simulations suggest an out-of-sample $R^2 \in [0.27, 0.78]$ is likely.

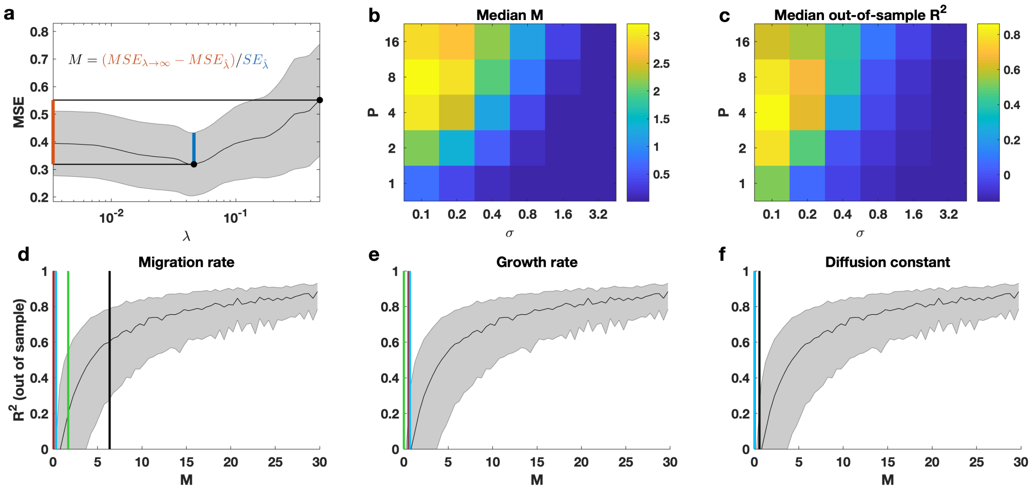


Figure S1: **Numerical investigation of LASSO regression demonstrates mutations have limited predictive power of migration phenotypes.** **(a)** Schematic demonstrating the $M$ statistic. $M$ measures the improvement of the model that minimizes cross-validation MSE relative to the trivial model with only an intercept term. This statistic is scaled by the estimated standard error of the MSE at its minimum to incorporate uncertainty in the minimum MSE estimate. **(b-c)** Results of surrogate data simulations at different values of $P$ (the number of true nonzero regression coefficients) and $\sigma$ (the standard deviation of the noise term). Median values of $M$ across $10^4$ simulations per $(P, \sigma)$ combination show that high $M$ is achieved in a high signal-to-noise regime, i.e., when $\sigma$ is sufficiently small and $P > 1$. Median out-of-sample $R^2$ values are also largest in this high signal-to-noise regime. **(d-f)** $M - R^2$ relationship from surrogate data with $M$ values from real data regressions overlaid. $M$ and $R^2$ values from all surrogate data simulations are combined, binned by $M$, and the quartiles of $R^2$ within the bins are shown as a function of $M$. The same quartile curves are shown in all three panels. Vertical lines indicate the $M$ values resulting from LASSO regressions on real data for adaptation in migration rate, growth rate, and diffusion constant **d,e,f** respectively, where the colors indicate the assay condition (black: mannose, red: melibiose, green: N-acetylglucosamine, blue: galactose). Comparison between the numerical investigation and the $M$ values from real data regressions suggests that mutation candidacy very likely does not predict migration-related phenotypic parameters reliably (predicted out-of-sample $R^2 \sim 0$), with the exception of migration rate adaptation assayed in mannose.

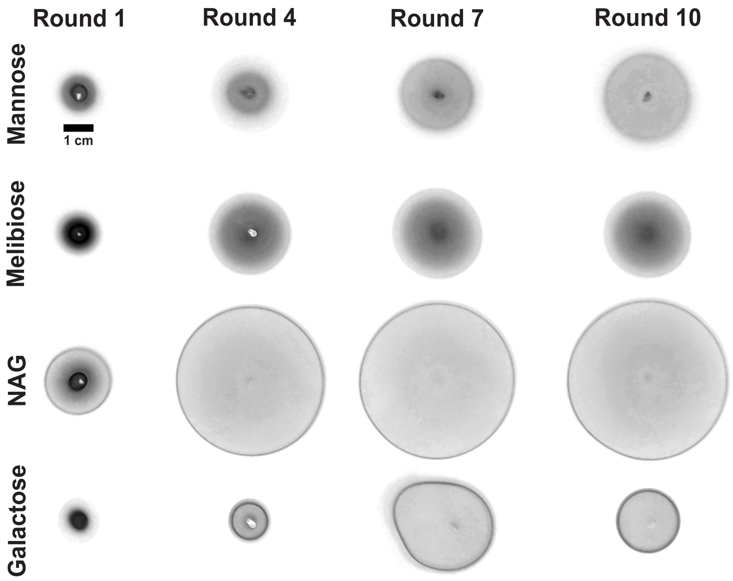


Figure S2: ***E. coli* colonies throughout selection experiment.** Example images of expanded colonies after 24 hours of migration at rounds 1, 4, 7 and 10 of the selection experiments described in the main text. 1 cm scale bar applies to all images. Darker regions correspond to higher cell density. Grayscale images were background subtracted, inverted, and had their dynamic range adjusted for better contrast. The asymmetry observed in galactose round 7 is due to inhomogeneity in the soft agar plate.

| Strain | Arabinose | Dextrose | Fructose | Lactose | Maltose | Rhamnose | Sorbitol |
|---|---|---|---|---|---|---|---|
| mannose 10A | $2.8 \pm 0.18$ | $2.1 \pm 0.14$ | $1.7 \pm 0.02$ | $1.6 \pm 0.11$ | $2.0 \pm 0.10$ | $1.7 \pm 0.30$ | $2.0 \pm 0.03$ |
| mannose 10B | $2.3 \pm 0.15$ | $2.2 \pm 0.03$ | $1.8 \pm 0.02$ | $1.6 \pm 0.06$ | $2.1 \pm 0.003$ | $1.2 \pm 0.12$ | $1.9 \pm 0.08$ |
| melibiose 10A | $2.1 \pm 0.18$ | $2.1 \pm 0.21$ | $1.9 \pm 0.20$ | $1.5 \pm 0.05$ | $1.7 \pm 0.01$ | $1.5 \pm 0.31$ | $1.9 \pm 0.05$ |
| melibiose 10B | $2.4 \pm 0.39$ | $1.9 \pm 0.01$ | $2.2 \pm 0.005$ | $1.5 \pm 0.04$ | $1.7 \pm 0.01$ | $2.3 \pm 0.29$ | $1.9 \pm 0.13$ |
| NAG 10A | $3.5 \pm 0.33$ | $3.0 \pm 0.09$ | $2.7 \pm 0.18$ | $1.7 \pm 0.14$ | $2.1 \pm 0.06$ | $2.3 \pm 0.08$ | $3.1 \pm 0.002$ |
| NAG 10B | $3.7 \pm 0.51$ | $3.0 \pm 0.27$ | $3.0 \pm 0.32$ | $1.9 \pm 0.01$ | $2.0 \pm 0.03$ | $2.6 \pm 0.04$ | $3.0 \pm 0.34$ |
| galactose 10A | $2.6 \pm 0.44$ | $2.5 \pm 0.001$ | $2.6 \pm 0.12$ | $1.6 \pm 0.02$ | $1.8 \pm 0.03$ | $2.4 \pm 0.41$ | $2.4 \pm 0.05$ |
| galactose 10B | $1.9 \pm 0.22$ | $1.9 \pm 0.04$ | $2.0 \pm 0.12$ | $1.2 \pm 0.04$ | $1.5 \pm 0.05$ | $2.0 \pm 0.04$ | $1.8 \pm 0.01$ |

Table S1: **Nutrient generality extends to a variety of other sugars.** We assayed the migration rates of the ancestor as well as two evolved strains isolated after 10 rounds from each selection condition in a variety of sugars. Rates are presented as fold change compared to the founder's migration rate in the same condition, mean $\pm$ standard deviation of two replicate plates for each strain in each condition. The founder has a migration rate of $0.026 \pm 0.001$, $0.062 \pm 0.005$, $0.034 \pm 0.004$, $0.069 \pm 0.002$, $0.069 \pm 0.005$, $0.027 \pm 0.002$, $0.047 \pm 0.004$ cm $\mathrm{h}^{-1}$ in arabinose, dextrose, fructose, lactose, maltose, rhamnose and sorbitol, respectively, mean $\pm$ standard deviation of two replicate plates in each condition.

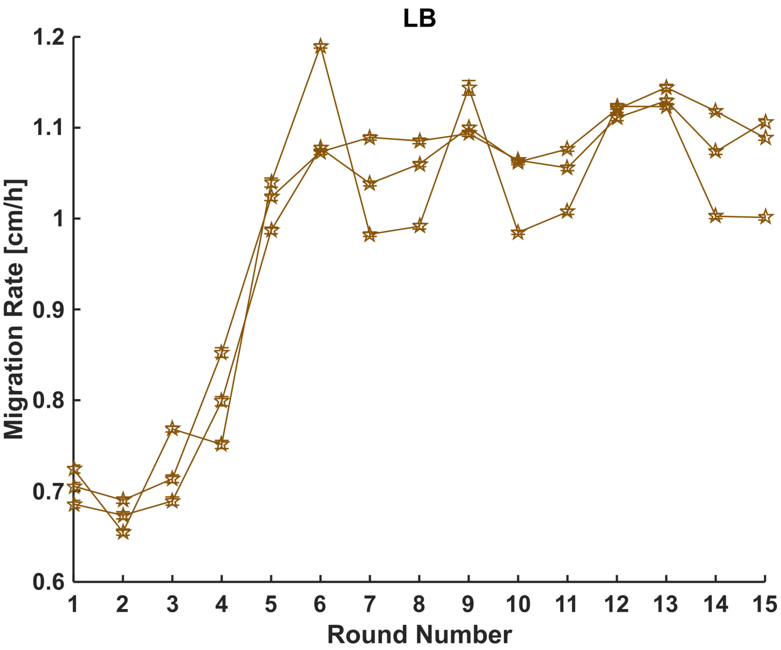


Figure S3: **Repeated selection enhances *E. coli* migration through soft agar in rich medium.** Migration rates as a function of round of selection for three replicate experiments performed in lysogeny broth (LB) rich medium. Selection and migration rate measurement were performed as described in main text methods, with the following exceptions: We used 15 cm petri dishes (containing LB with 0.2 % w/v agar) and selection was performed every 8 hours due to the fast migration rates in this condition. Seed cultures were grown overnight in 5 mL LB and time-lapse imaging was performed every minute. After imaging, four 50 µL samples were removed from the outermost edge of the expanding colony. During image analysis, front location was determined by locating peaks in the radial density profiles. These experiments were carried out to 15 rounds, however, the strains isolated after only 10 rounds were used for the comparison with 10-round minimal medium evolved strains described in the main text and presented below.

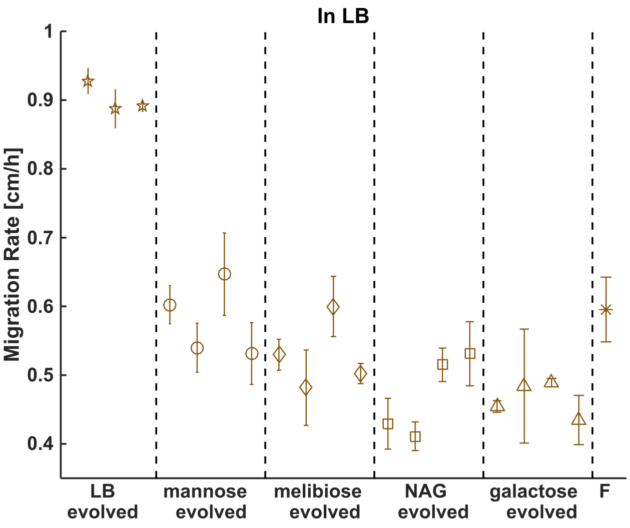


Figure S4: **Nutrient generality does not extend to rich medium.** The 16 strains isolated after 10 rounds of selection (four from each nutrient condition, main text Figure 1) were assayed for enhanced migration rate in LB rich medium. Migration rates of these strains are presented as mean ± standard deviation of two replicate plates. For comparison, we also measured migration rates of the founding strain (F) as well as three strains isolated after 10 rounds of selection in LB (figure S3). Migration rates of these strains are presented as mean ± standard deviation of four replicate plates. Migration rate assays were conducted as described in main text methods, with the following exceptions: We used 10 cm petri dishes containing LB with 0.2 % w/v agar. Seed cultures were grown overnight in 5 mL LB and time-lapse imaging was performed every two minutes for eight hours. Only the first five hours were analyzed, since the LB-evolved strains reach the boundary of a 10 cm plate around this time. During image analysis, front location was determined by locating peaks in the radial density profiles.

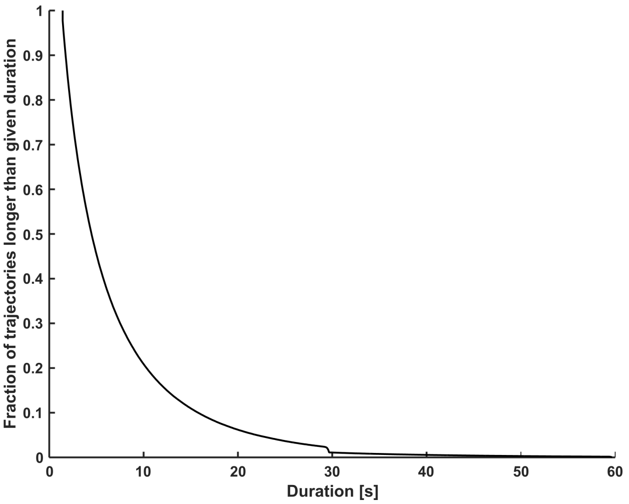


Figure S5: **Most trajectories do not extend past the time interval used for fitting MSD.** Complementary cumulative distribution function of trajectory duration observed in all single-cell tracking experiments. To measure diffusion constant we fit mean-squared displacement over the interval from 1 to 6 seconds into the MSD trace (Figure 4a, main text).

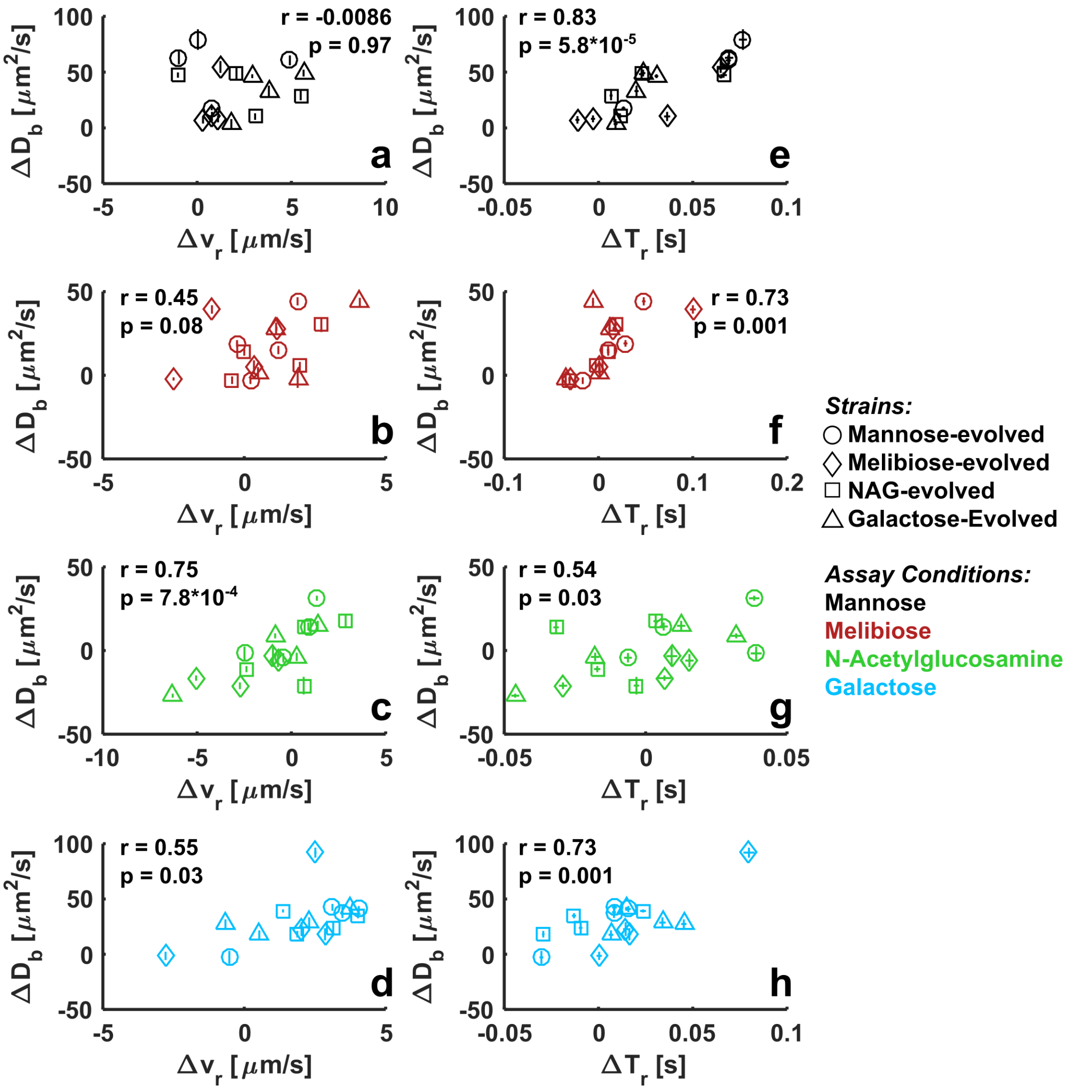


Figure S6: **Plasticity in $\Delta D_b$ extends to run-tumble statistics.** For each single-cell tracking experiment presented in main text Figure 4c, we detected runs and tumbles using the Hidden Markov Model classifier in Pytaxis. For each detected run, we compute its duration and average speed. We average over all runs in an experiment to obtain mean run speed $v_r$ and duration $T_r$ of each strain in each condition. The founder has a run speed of $16.1 \pm 0.5, 17.4 \pm 0.3, 16.5 \pm 0.7$ and $16.8 \pm 0.4$ $\mu m\, s^{-1}$ and a run duration of $0.29 \pm 0.002, 0.32 \pm 0.02, 0.31 \pm 0.03$ and $0.31 \pm 0.007$ seconds in mannose, melibiose, NAG and galactose respectively, mean $\pm$ standard deviation of two replicate experiments. We subtract these values from the evolved strains to obtain $\Delta v_r$ and $\Delta T_r$ for each evolved strain in each condition. $\Delta D_b$ are reproduced from main text Figure 4c. For each panel, we obtain a Pearson correlation coefficient and associated p-value. For strains measured in mannose and melibiose, we obtain significant ($p < 0.05$) correlations between $\Delta D_b$ and $\Delta T_r$, but not between $\Delta D_b$ and $\Delta v_r$. We conclude that strains measured in these conditions increase their diffusion constant by extending run duration. For strains measured in N-acetylglucosamine and galactose, we obtain significant correlations between $\Delta D_b$ and both $\Delta v_r$ and $\Delta T_r$. However, changes in $D_b$ were correlated more strongly with run speed than run duration in N-acetylglucosamine and the converse in galactose.

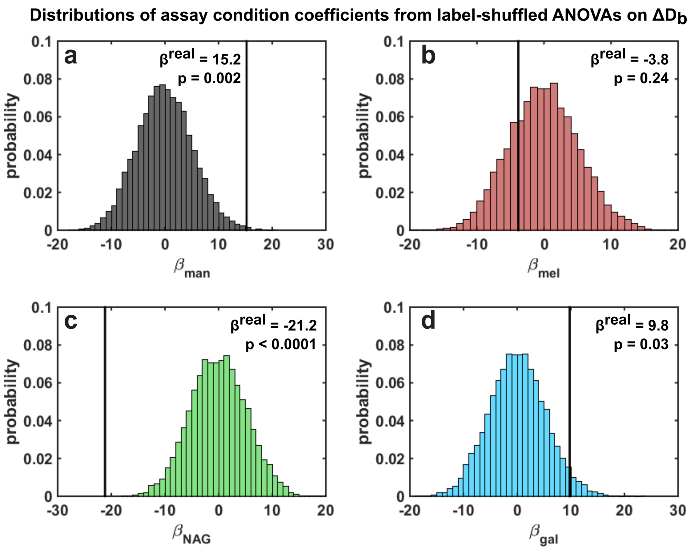


Figure S7: **Three assay conditions have significant effects on diffusion adaptation.** Results of bootstrapping approach used to determine which assay conditions show significant ($p < 0.05$) departures from the global mean in $\Delta D_b$, and in which direction. ANOVA coefficients from the properly-labeled data set are presented as $\beta^{real}$ (vertical black lines). p-values are computed by determining the fraction of corresponding coefficients from label-shuffled ANOVAs that are higher or lower than $\beta^{real}$, depending on its sign. $p < 0.0001$ indicates none of the 10 000 ANOVAs on label-shuffled data had a more negative coefficient than $\beta^{real}_{NAG}$ See main text methods for details.

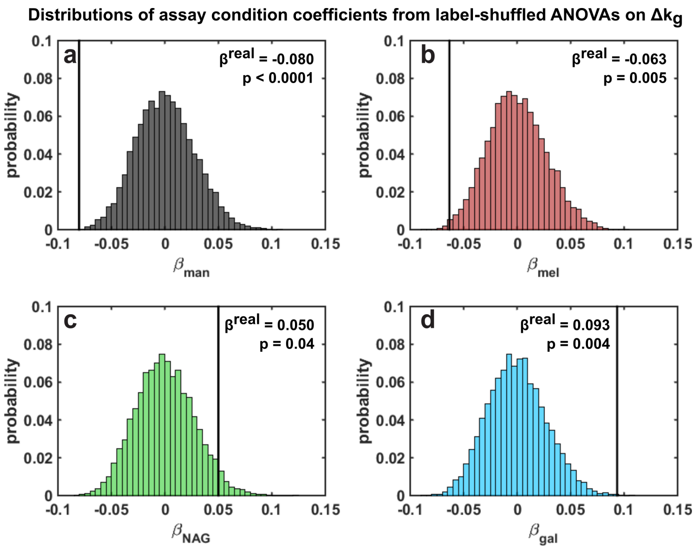


Figure S8: **All four assay conditions have significant effects on growth rate adaptation.** Results of bootstrapping approach used to determine which assay conditions show significant ($p < 0.05$) departures from the global mean in $\Delta k_g$, and in which direction. ANOVA coefficients from the properly-labeled data set are presented as $\beta^{real}$ (vertical black lines). p-values are computed by determining the fraction of corresponding coefficients from label-shuffled ANOVAs that are higher or lower than $\beta^{real}$, depending on its sign. $p < 0.0001$ indicates none of the 10 000 ANOVAs on label-shuffled data had a more negative coefficient than $\beta^{real}_{man}$. See main text methods for details.

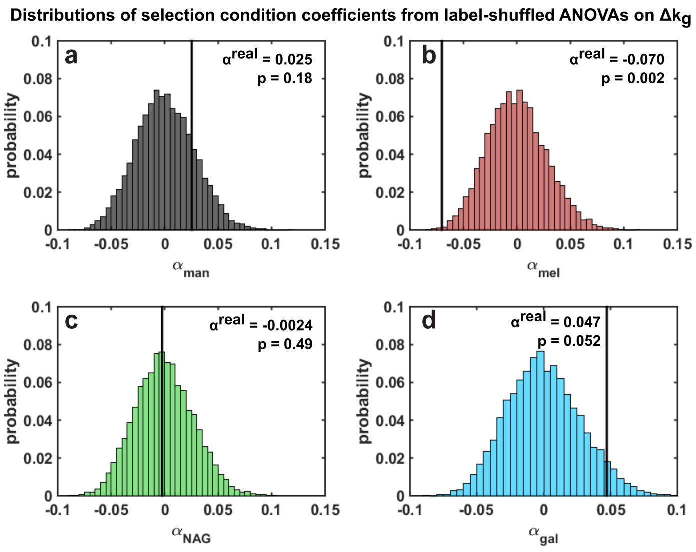


Figure S9: **Melibiose-evolved strains have below-average growth rate adaptation.** Results of bootstrapping approach used to determine which selection conditions show significant ($p < 0.05$) departures from the global mean in $\Delta k_g$, and in which direction. ANOVA coefficients from the properly-labeled dataset are presented as $\alpha^{real}$ (vertical black lines). p-values are computed by determining the fraction of corresponding coefficients from label-shuffled ANOVAs that are higher or lower than $\alpha^{real}$, depending on its sign. See main text methods for details.

| **Interaction Coefficient** | $(\alpha\beta)^{real}$ | **p-value** |
|---|---|---|
| SelectionCond=man x AssayCond=man | 0.052 | 0.14 |
| SelectionCond=man x AssayCond=mel | -0.005 | 0.47 |
| SelectionCond=man x AssayCond=nag | 0.005 | 0.44 |
| SelectionCond=man x AssayCond=gal | -0.052 | 0.14 |
| SelectionCond=mel x AssayCond=man | 0.011 | 0.39 |
| SelectionCond=mel x AssayCond=mel | 0.042 | 0.19 |
| SelectionCond=mel x AssayCond=nag | -0.015 | 0.39 |
| SelectionCond=mel x AssayCond=gal | -0.039 | 0.21 |
| SelectionCond=nag x AssayCond=man | -0.028 | 0.28 |
| SelectionCond=nag x AssayCond=mel | 0.008 | 0.42 |
| SelectionCond=nag x AssayCond=nag | 0.089 | 0.04 |
| SelectionCond=nag x AssayCond=gal | -0.069 | 0.06 |
| SelectionCond=gal x AssayCond=man | -0.035 | 0.23 |
| SelectionCond=gal x AssayCond=mel | -0.045 | 0.18 |
| SelectionCond=gal x AssayCond=nag | -0.080 | 0.04 |
| SelectionCond=gal x AssayCond=gal | 0.160 | 0.001 |

Table S2: **NAG-evolved and galactose-evolved strains have a 'home field advantage' in growth rate adaptation.** Results of bootstrapping approach used to determine which selection x assay interaction terms show significant ($p < 0.05$) departures from the global mean in $\Delta k_g$, and in which direction. ANOVA coefficients from the properly-labeled data set are presented as $(\alpha\beta)^{real}$. p-values are computed by determining the fraction of corresponding coefficients from label-shuffled ANOVAs that are higher or lower than $(\alpha\beta)^{real}$, depending on its sign. See main text methods for details.

| Targets | Mutations observed | Present in these strains |
|---|---|---|
| glyA | H165H | man10A, man10C, man10D, nag10A, nag10C, nag10D, gal10B |
| yegH | R335R | man10B, mel10B, gal10A |
| rpoB$^{12}$, rpoC$^{3}$ | P552L$^{1}$, I569L$^{2}$, +9bp$^{3}$ | man10D$^{1}$, nag10A$^{2}$, nag10C$^{3}$ |
| rssB | A280T | mel10A, mel10D, gal10B, gal10C, gal10D |
| mepS$^{1245}$<br>lpxT→ / →mepS$^{3}$ | IS5(–)+4bp$^{1}$, E171*$^{2}$<br>IS1(+)+8bp$^{34}$, IS1(+)+9bp$^{5}$ | mel10A$^{1}$, mel10B$^{2}$, nag10B$^{3}$<br>nag10C$^{4}$, nag10D$^{5}$ |
| glxK | P210L | mel10B, mel10C, mel10D |
| yeaR | Δ1::IS186(–)+6bp::Δ1 | mel10B, gal10B |
| frdA | G393V | mel10B, gal10B |
| envZ | Δ36bp$^{1}$, Δ1bp$^{2}$ | nag10A$^{1}$, nag10D$^{2}$ |
| rph$^{13}$<br>pyrE← / ←rph$^{2}$ | Δ1bp$^{1}$, A→G$^{2}$<br>+A$^{3}$ | nag10B$^{1}$, gal10A$^{2}$, gal10D$^{3}$ |
| nagA | Δ1bp$^{13}$, A→C$^{2}$<br>Δ10bp$^{4}$ | gal10A$^{12}$, gal10B$^{34}$ |
| metK→ / →galP | C→A$^{1}$, G→T$^{2}$ | gal10C$^{1}$, gal10D$^{2}$ |
| yghG | E30* | mel10C |
| mokB← / →trg | G→T | mel10D |
| wzzE | IS1(–)+9bp | nag10B |
| yffR→ / →yffS | C→A | nag10D |
| osmC | D90N | gal10B |
| yncE→ / ←ansP | C→A | gal10C |
| yggI | G159G | gal10C |
| ligB | A142V | gal10D |
| nrfG→ / →gltP | G→T | gal10D |

Table S3: **The set of mutations shared between strains with different evolutionary histories.** Novel mutations present at frequencies of 20 % or greater in the 16 evovled strains presented in main text Figure 2. Whole-genome sequencing and analysis was performed as described in main text methods with an average coverage of $59.8 \pm 11.2$ (mean $\pm$ standard deviation across strains). We group mutations by target since some genes exhibit different mutations across strains and since some strains have mutations in intergenic regions adjacent to genes affected in other strains. For these cases, superscripts indicate which strains had which mutations, and whether they occurred in the coding region or the intergenic space (that is, superscripts are specific to each row of the table where they appear). This target-level grouping was used for the candidacy matrix used in the LASSO regressions. Strains are designated by their selection condition (mannose, melibiose, N-acetylglucosamine, galactose), rounds of selection (10 for these strains) and replicate (A,B,C,D, since four independent lineages were sequenced from each selection condition). Notation convention for mutations is described in the *breseq* documentation (`http://barricklab.org/twiki/pub/Lab/ToolsBacterialGenomeResequencing/documentation`).
